# Towards edge processing of images from insect camera traps

**DOI:** 10.1101/2024.07.01.601488

**Authors:** Kim Bjerge, Henrik Karstoft, Toke T. Høye

**Affiliations:** Department of Electrical and Computer Engineering, Aarhus University, Aarhus C, Denmark; Department of Ecoscience, Aarhus University, Aarhus C, Denmark; Arctic Research Center, Aarhus University, Aarhus C, Denmark

**Keywords:** Biodiversity monitoring, Camera trap, Computer Vision, Edge processing, Insect classification, Tracking

## Abstract

Insects represent nearly half of all known multicellular species but knowledge about them lacks behind most vertebrate species. In part for this reason, they are often neglected in biodiversity conservation policies and practice. Computer vision tools, such as insect camera traps, for automated monitoring have the potential to revolutionize insect study and conservation. To further advance insect camera trapping and the analysis of their image data, effective image processing pipelines are needed. In this paper, we present a flexible and fast processing pipeline designed to analyse these recordings by detecting, tracking and classifying nocturnal insects in a broad taxonomy of 15 insect classes and resolution of individual moth species. A classifier with anomaly detection is proposed to filter dark, blurred, or partially visible insects that will be uncertain to classify correctly. A simple track-by-detection algorithm is proposed to track classified insects by incorporating feature embeddings, distance and area cost. We evaluated the computational speed and power performance of different edge computing devices (Raspberry Pi’s and NVIDIA Jetson Nano) and compared various time-lapse strategies with tracking. The minimum difference was found for 2-minute time-lapse intervals compared to tracking with 0.5 frames per second, however, for insects with fewer than one detection per night, the Pearson correlation decreases. Shifting from tracking to time-lapse monitoring would reduce the amount of recorded images and be able to perform edge processing of images in real-time on a camera trap with Raspberry Pi. The Jetson Nano is the most energy-efficient solution, capable of real-time tracking at nearly 0.5 fps. Our processing pipeline was applied to more than 5.7 million images recorded at 0.5 frames per second from 12 light camera traps during two full seasons located in diverse habitats, including bogs, heaths and forests.

## 1 INTRODUCTION

Insects make up the most diverse group of animals with more than a million described species and insects constitute 50% of the total animal biomass (Bar-On et al. 2018). Insects play vital roles in terrestrial ecosystems and have significant economic importance as agricultural pests, natural enemies and pollinators. Changes in insect abundance have cascading effects through the food web. suggesting that insects are a very relevant animal group for monitoring more effectively in the context of global change (Wagner et al. 2021). Conventional insect trapping techniques, as outlined by Montgomery et al. (2021), are labour intensive and in many cases insects are sacrificed in the process. Manual enumeration and taxonomic identification by human experts is also very labour intensive and often requires highly specialized knowledge.

Data on insect populations is notably sparse due to limited resources, the vast number of species and the high level of expertise required to study them (Didham et al. 2020). The advent of automated monitoring technologies, employing computer vision and deep learning, has brought about a revolution in insect studies (van Klink et al. 2022, Lima et al. 2020, Besson et al. 2022) in both real-time scenarios (Bjerge et al. 2021a, Sittinger et al. 2024, Ratnayake et al. 2021) and offline analysis of images from time-lapse (TL) cameras (Geissmann et al. 2022, Bjerge et al. 2023a). Automated insect camera traps, coupled with data-analysing algorithms rooted in computer vision and deep learning, could therefore serve as invaluable tools to monitor insect trends and elucidate the underlying drivers (Barlow and O’Neill 2020, Høye et al. 2021). Animal species recognition from camera traps is a well-established problem within the computer vision community (Oliver et al. 2023), with common challenges including poor lighting, occlusion, camouflage and blur (Beery et al. 2018). However, working with insects presents unique challenges that are not encountered with traditional camera trap systems designed for large animals. For example, while traditional camera trap images might occasionally capture a target species, nearly every image from an insect camera trap contains insects. This is especially true during nights of high activity, where hundreds of nocturnal insects can be visible in a single image.

Nocturnal insects are difficult to monitor, however, camera-based light traps (Bjerge et al. 2021b, Korsch, D., Bodesheim, P., Denzler, J. 2021) and advancement of standardized hardware and framework for image-based monitoring of nocturnal insects (Roy et al. 2024) paves the way for increased temporal coverage and resolution in insect monitoring. Automated monitoring of moths has been evaluated by comparing traditional lethal methods with light-based camera traps (Möglich et al. 2023). This first proof of concept has demonstrated that automated moth traps captures phenological patterns just as well as conventional, lethal traps.

Camera trapping methods based on time-lapse recordings can generate millions of images, especially when using sampling intervals of seconds or a few minutes. However, as these tools become more widely applied, they are likely to generate large amounts (terabytes to petabytes) of image data per year and storing all the data may not be feasible or even sensible. It is possible that a reduced frame rate will yield comparable results, but rare taxa are less likely to be detected as the frame rate is reduced. An alternative approach is to implement edge computing, where image processing is performed directly on the recording camera device. In this setup, only the processed data and, optionally, a subset of raw images are stored. Edge computing facilitates real-time monitoring, allowing daily uploads of insect taxa abundance statistics when internet access is available. However, edge computing requires significant computational power, which increases the cost of the camera system.

### 1.1 Contribution

In this work, we propose a flexible and fast processing pipeline to analyse image recordings from insect camera traps by detecting, tracking and classifying nocturnal insects at the broad taxonomic ranks such as order, suborder, family and at the species level for moths. We demonstrate the efficacy of the proposed pipeline by evaluating its speed performance on three different edge computing devices, including Raspberry Pi 4, Raspberry Pi 5 and NVIDIA Jetson Nano. Our pipeline supports multiple time-lapse strategies and real-time tracking. These strategies are evaluated to ensure they provide comparable measurements of activity dynamics over time. We apply the pipeline to image data recorded with 12 insect camera traps fitted with UV light to attract nocturnal insects. The dataset includes recordings from >3000 nights across two years. The statistics of recorded images, detected and tracked insects from this study are presented in this paper.

In summary, our objectives for this study are the following:

- Propose a deep learning pipeline to measure temporal abundance for taxa of nocturnal insects.
- Classify all images of insects into broad taxonomic groups with anomaly detection and images of Lepidoptera to species.
- Evaluate four different computing platforms including edge devices with respect to processing time and energy consumption.
- Compare time-lapse sampling with real-time tracking of individual insects.
- Connect the pipeline to insect ecology and conservation by demonstrating the proposed image processing pipeline in field-collected data.

## 2 MATERIALS AND METHOD

### 2.1 Data collection

Automated light traps with cameras were constructed with standard components consisting of a Raspberry Pi 4, Brio Camera (Logitech 2021), power controller, UV light and light ring as proposed by Bjerge et al. (2021b). A solid-state drive (500 GB) was connected to the Raspberry Pi to store the captured images. The mechanical design was improved and the background light table was replaced with a plastic plate covered with a white fabric shown in Figure 1.

**FIGURE 1.**
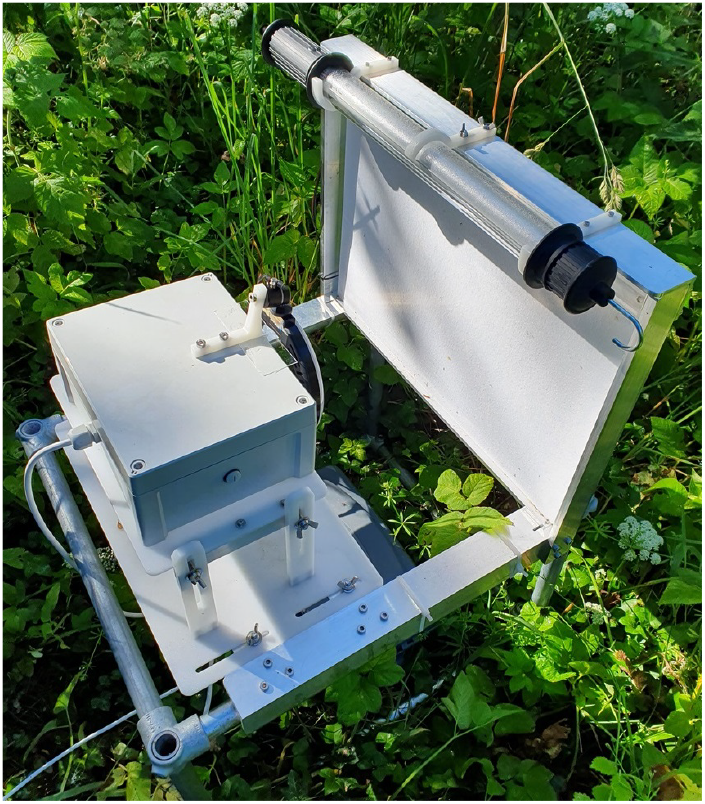
Camera trap with UV light to attract and monitor nocturnal insect.

12 camera traps were placed at three different locations in Denmark during the 2022 and 2023 summer seasons. These locations included a variety of habitats, such as bogs, heaths and forests. Three nature areas managed by the Aage V. Jensen Naturfond were selected for sampling. Lille Vildmose, Ovstrup Hede and Søholt Storskov ^†^. Within each area, four traps were deployed, spaced 1–10 km apart. Four of the traps were powered by solar panels, charge regulators and batteries (12V), all other traps were supplied through mains power (220V). The camera traps were activated in the period from 11 p.m. to 3 a.m. each night. We restricted sampling to this period each night to ensure that power from the battery and solar panel was available throughout the entire insect activity season in Denmark from April to the end of October, i.e. even when the solar angle is fairly low at the sites. We turned off the traps at 3 a.m. to ensure that insects would have sufficient time to leave the trap before insectivorous birds would become active in the morning.

A motion program (Motion 2021) running on the Raspberry Pi 4 was installed to capture a sequence of images whenever a movement was detected in the camera view. The maximum frame rate was limited to 0.5 fps. On warm summer nights with a high level of insect activity, more than 6,000 images were captured per night. In 2022 and 2023 more than 5 million images with a pixel size of 3840 × 2160 (11 pixels/*mm*) were recorded. As a supplement to the motion recorded images, a time-lapse approach was used to save an image every 10 minutes independent of insect activity.

### 2.2 Processing pipeline

For insect monitoring, Multiple Object Tracking (MOT) would be relevant, especially for fast video recording. MOT uses Computer Vision to estimate trajectories for objects of interest presented in a sequence of images, especially videos with high frame rates. Most MOT methods require annotated tracking datasets, which can be challenging to create.

We aimed to create a flexible pipeline that can be used for both processing with and without tracking depending on the chosen time-lapse sampling interval. We chose the track-by-detection (TBD) approach since it is flexible and can be realized without any annotated tracking dataset. The proposed processing pipeline is shown in Figure 2. The pipeline is designed to prioritize flexibility above efficiency.

**FIGURE 2.**
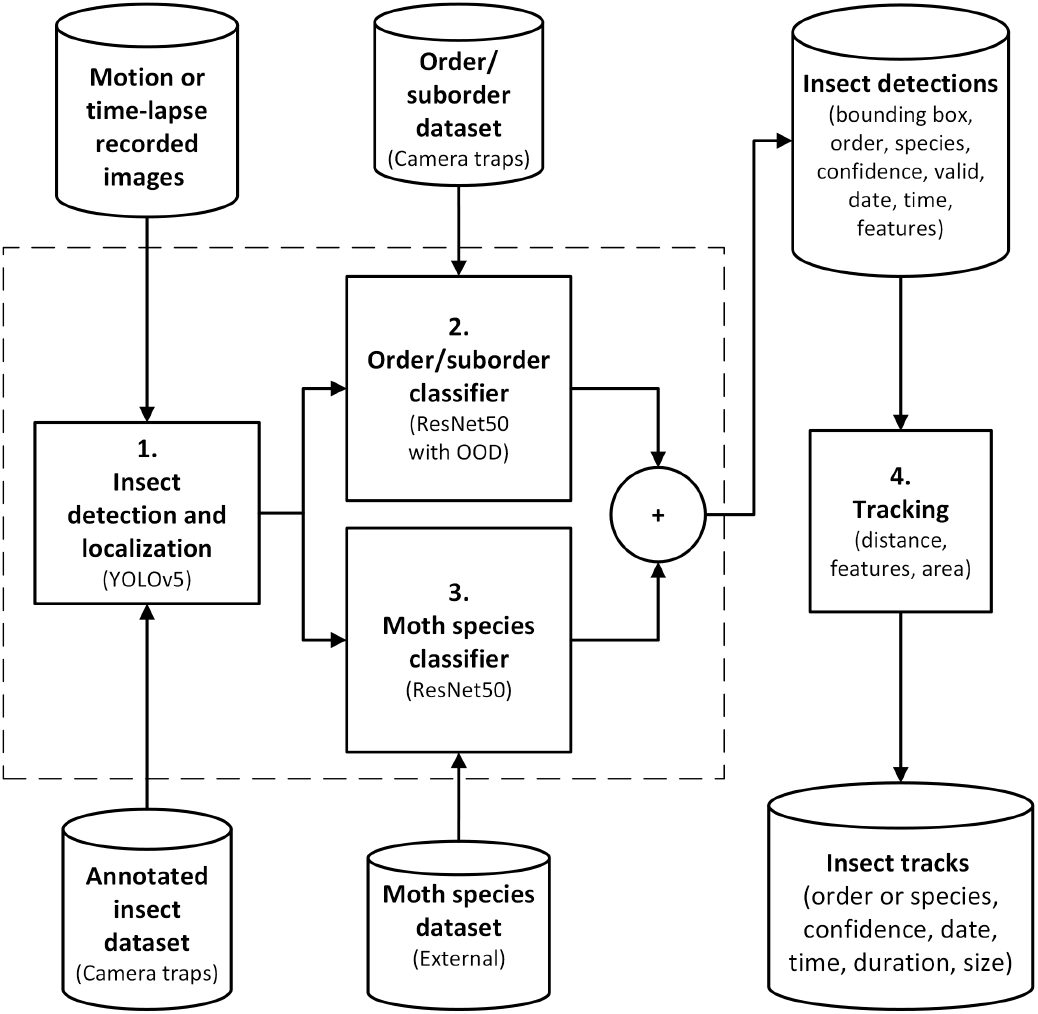
Processing pipeline to localize, classify and track insects from motion or time-lapse recorded camera trap images. In steps 1. and 2., the detected insects are classified to the level of broad taxonomic groups parallel with moth species classification. Classification results are concatenated “+” to provide insect information for the final tracking step performed on motion triggered images recorded with a high sampling rate of 0.5 fps.

**The first step** in our pipeline is to detect insects. To perform this step, we first annotated a training dataset of insects in the images collected from our camera traps. We then trained a model to detect insects of interest while ignoring dirt and small or blurry insects. In the future, this step could be replaced with a more generic detector trained on images from various backgrounds with insects.

**The second and third step** classifies the detected insects using two separate models. One classifies all insects into broad taxonomic groups and the other classifies moths to the species level. The two classification models can be executed in parallel.

The broad taxon classifier is trained on the camera trap data. Here, we have sorted the insects into order, suborders and families based on the content of the recorded images. We have incorporated the anomaly detector presented by (Bjerge et al. 2023b) into the classifier to filter insects which have class scores that are outside the distribution of the created dataset of broad taxonomic groups. These outliers could be partly visible or blurry insects, or they could be representatives of unseen classes of insects or other animals.

**The third step** implements the moth species classifier trained on external data from the Global Biodiversity Information Facility (GBIF) with all moth species known to be present in the region where the trap is located. Here, we have used the moth species classifier published by Rolnick, David; Bundsen, Michael; Jain, Aditya; Cunha (2023) trained for moth species found in Denmark and the UK.

The output from the insect detector, broad taxon classifier and moth species classifier is a list of insect detections with additional information about the trap, image, bounding box coordinates, confidence, anomaly, date, time and embedding features. **The fourth tracking step** is based on track-by-detection by using the bounding boxed and embedding features to create a final list of insect tracks with information about predicted insect taxon, species, confidence, size, date, arrival time and duration seen by the camera.

The source code for the pipeline is available on Github^‡^. Each step in the pipeline is described below, with a focus on the contributions for anomaly detection and the simple flexible tracking of insects.

### 2.3 Insect detection and localization

Deep learning image object detection methods rely solely on spatial image information to extract features and detect regions of objects in the image. You-only-look-once (Redmon et al. 2016) (YOLO) is a one-stage object detector and one of the fastest object detectors important for processing millions of images or deployed on edge computers. In our work, YOLOv5 (Glenn Jocher 2020) with CSPDarknet53 as the backbone was evaluated.

In the paper Bjerge et al. (2023a) different YOLOv5 architectures are evaluated, finding that YOLOv5m6 with 35.7 million parameters is the optimal model to detect and classify small insect species. To improve performance and speed up training, YOLOv5m6 is pre-trained on the Common Objects in Context (COCO) dataset (Lin et al. 2015) that contains more than 330,000 images of 80 different categories of objects. In this work, we have fine-tuned YOLOv5m6 and YOLOv5s6 on the dataset described in Section 2.7.

### 2.4 Broad taxon classifier with anomaly detection

The images were cropped and resized to 128 × 128 pixels, which matches the dimensions used for the moth species classifier. The training on the datasets was performed using data augmentation, including image scaling, horizontal and vertical flip and adding color jitter for brightness, contrast and saturation. We selected a batch size of 256 for training our models, since it is faster to update and results in less noise than smaller batch sizes. The Adam optimizer with a fixed learning rate of 1.0 · 10^−4^ was chosen based on previously published experiments (Bjerge et al. 2023b). We have trained ResNet50v2 (He et al. 2016) to classify insects according to broad taxonomic groups defined by the 16 classes as described in Section 2.7. ResNet50v2 was fine-tuned using pre-trained weights from ImageNet (Russakovsky et al. 2015).

#### 2.4.1 Out-of-distribution detection

The methodology of out-of-distribution detection (Bulusu et al. 2020) and threshold-based anomaly tagging is employed to identify instances of “anomalies” such as uncertain classifications. In our application, these instances may manifest as debris, obscured or partially visible insects, or those exhibiting blurriness, characteristics that are not represented in the insect taxon training dataset.

Often, softmax is the last layer in a classification neural network, where the maximum value determines the predicted class. Here, we instead analyse the output distribution without the softmax layer to determine the anomalies and predict the classes. The distribution of the output *x* for each predicted class *j*^*th*^ follows a normal distribution 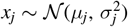.

An example of the output distribution is shown in Figure 3 which is generated on the sample training dataset (Diptera Brachycera) for corrected classified inputs. If the output value *x*_*j*_ is below a threshold of *th* = *µ* – 2.5*σ*, we label the input as anomaly. Consequently, when new unknown inputs are presented for the trained network and the output lies below the threshold, it will be classified as an “uncertain” prediction. The threshold is set to ensure that fewer than 1% of the correctly classified inputs are discarded. However, thresholds between *µ* – 2.0*σ* and *µ* – 3.0*σ* can also be selected, depending on the desired strictness of the anomaly detector.

**FIGURE 3.**
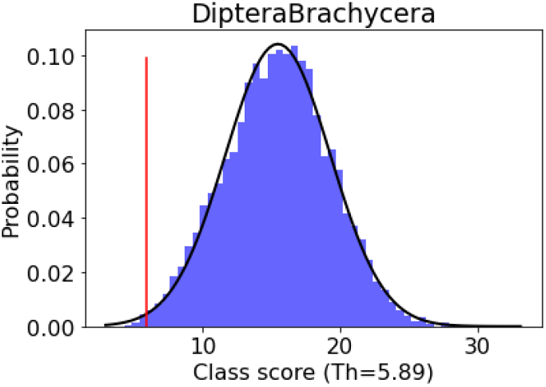
Probability density function for the output scores for Diptera Brachycera and the chosen threshold for uncertain anomalies.

Finally, the output scores *x*_*j*_ are assigned a probability *F*(*x*_*j*_) by estimating the cumulative distribution function as the integral of the probability density function given by.

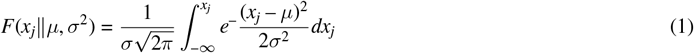

### 2.5 Moth species classifier

We use the moth species classifier from the companion code base of the AMI data (Jain et al. 2025, Rolnick, David; Bundsen, Michael; Jain, Aditya; Cunha 2023), trained on GBIF data that encompass 2,530 moth species found in the United Kingdom (UK) and Denmark. The model is tested on a dataset of moths recorded with AMI traps in Denmark and UK (Jain et al. 2025) with a F1-score of 0.784. All models within the AMI data companion code base are trained using the ResNet50 architecture. We anticipate that new classification models, covering diverse regions worldwide, will become available in the future, further enhancing the applicability and scope of moth species classification.

### 2.6 Tracking

Our tracking algorithm was extended by comparing feature embeddings from the broad taxon classifier for the tracking algorithm proposed by Bjerge et al. (2021a). The Hungarian Algorithm is the chosen method for finding the optimal assignment for a given cost matrix. In this application, the cost matrix should represent how likely it was that an insect in the previous image had moved to a given position in the current image. The cost function was defined as a weighted cost of embeddings similarity, distance and area of matching bounding boxes in the previous and current images. The Euclidean distance *D* between the center position (*x, y*) in the two images was calculated as follows.

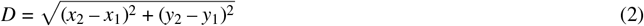

This distance was normalized according to the diagonal of the image *I*:

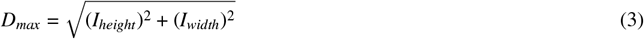

The area cost was defined as the cost between the area *A* of bounding boxes:

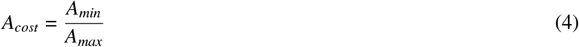

The similarity of feature embeddings was defined as the cosine similarity between embeddings ***E*** of classified insects:

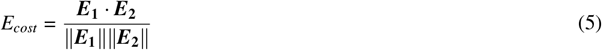

A final cost function in Equation (6) was defined with a weighted cost of distance *W*_*d*_, embeddings *W*_*e*_ and weighted cost of area *W*_*a*_.

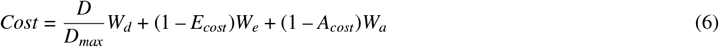

A cost threshold was established to determine whether successive insect detections should be associated. Subsequently, a track was established, stipulating a minimum of two detections per track. For each track, information such as the start date and time, duration, number of detections and average size was recorded. Of particular importance was the recording of the predominant insect taxon or moth species, along with the accuracy of its classification. Ultimately, a track was considered valid when >50% of the detections corresponded to the predominant classification and comprised at least three detections or had a duration of more than four seconds.

### 2.7 Datasets

The time-lapse recorded images were annotated to generate two distinct datasets aimed at facilitating insect localization followed by classification into broad taxonomic groups. By reviewing the detected insects, we identified 10 orders of insects (Coleoptera, Diptera, Ephemeroptera, Hemiptera, Hymenoptera, Lepidoptera, Neuroptera and Trichoptera) and arachnids (Araneae and Opiliones) frequently occurring in the dataset. For Diptera, Hymenoptera and Lepidoptera, it was also clear that the image quality allowed us to identify morphologically distinct taxonomic groups below the taxonomic level of order. Our aim was to balance the taxonomic resolution of the broad taxon classifier with the amount of training data per class that could be identified with a reasonable time investment. The arbitrary but pragmatic separation of macro and micro Lepidoptera was performed by grouping species of Lepidoptera at the family level. For more detailed future ecological analyses, it would be relevant to split the insect and arachnid taxa into further subgroups. This two-step strategy was adopted to manage the challenge of curating well-balanced datasets that include annotated insect taxa throughout the image dataset. This approach not only addresses the complexity of dataset creation, but also enables flexibility in the processing pipeline.

For detection, 777 images were selected and annotated, skipping very small and blurry insects from being annotated with bounding boxes. The 777 images were carefully selected to represent a diverse range of scenarios, including images from various camera traps that feature different insect species. The selection also included challenging cases, such as images with spider webs, dirt, blurry insects and insects that obscuring the camera lens. The complete dataset is shown in Table 1.

**TABLE 1.**
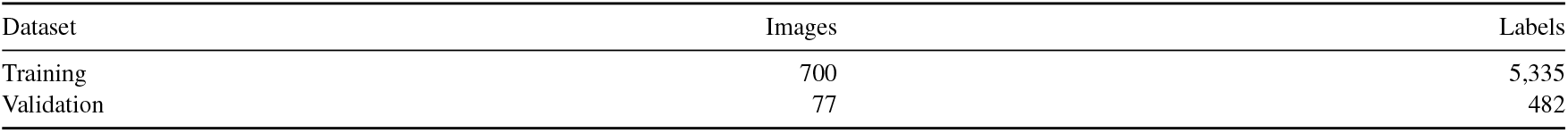
Dataset for detection and localization.

| Dataset | Images | Labels |
| --- | --- | --- |
| Training | 700 | 5,335 |
| Validation | 77 | 482 |

Example images of these taxa are provided in Figure 4. The resulting dataset, presented in Table 2, is organized according to the hierarchical rank of taxonomy. Furthermore, recognizing the inadvertent presence of vegetation, such as leaves and flowers, within the images, an additional class dedicated to vegetation was incorporated into the dataset to ensure comprehensive coverage of the observed objects. The dataset was split in 80% for training and 20% for validation of the classifier.

**TABLE 2.** Dataset of image samples for broad taxon classification collected during 2022 and 2023 from time-lapse images with 10-minute intervals.

| Taxa in Order [Suborder] | Taxon rank | Total samples | Validation (20%) |
| --- | --- | --- | --- |
| Araneae | Order | 2,037 | 408 |
| Coleoptera | Order | 2,384 | 477 |
| Diptera Brachycera | Suborder | 12,303 | 2,461 |
| Diptera Nematocera | Suborder | 26,890 | 5,378 |
| Diptera Tipulidae | Suborder | 1,216 | 244 |
| Diptera Trichoceridae | Suborder | 1,664 | 333 |
| Ephemeroptera | Order | 10,147 | 2,030 |
| Hemiptera | Order | 4,897 | 980 |
| Hymenoptera Other | Suborder | 1,661 | 333 |
| Hymenoptera Vespidae | Family | 529 | 106 |
| Lepidoptera Macro | Unranked | 18,675 | 3,735 |
| Lepidoptera Micro | Unranked | 28,141 | 5,629 |
| Neuroptera | Order | 1,106 | 222 |
| Opiliones | Order | 615 | 123 |
| Trichoptera | Order | 11,978 | 2,396 |
| Vegetation | Unranked | 892 | 179 |
| Total |  | 125,135 | 25,035 |

**FIGURE 4.**
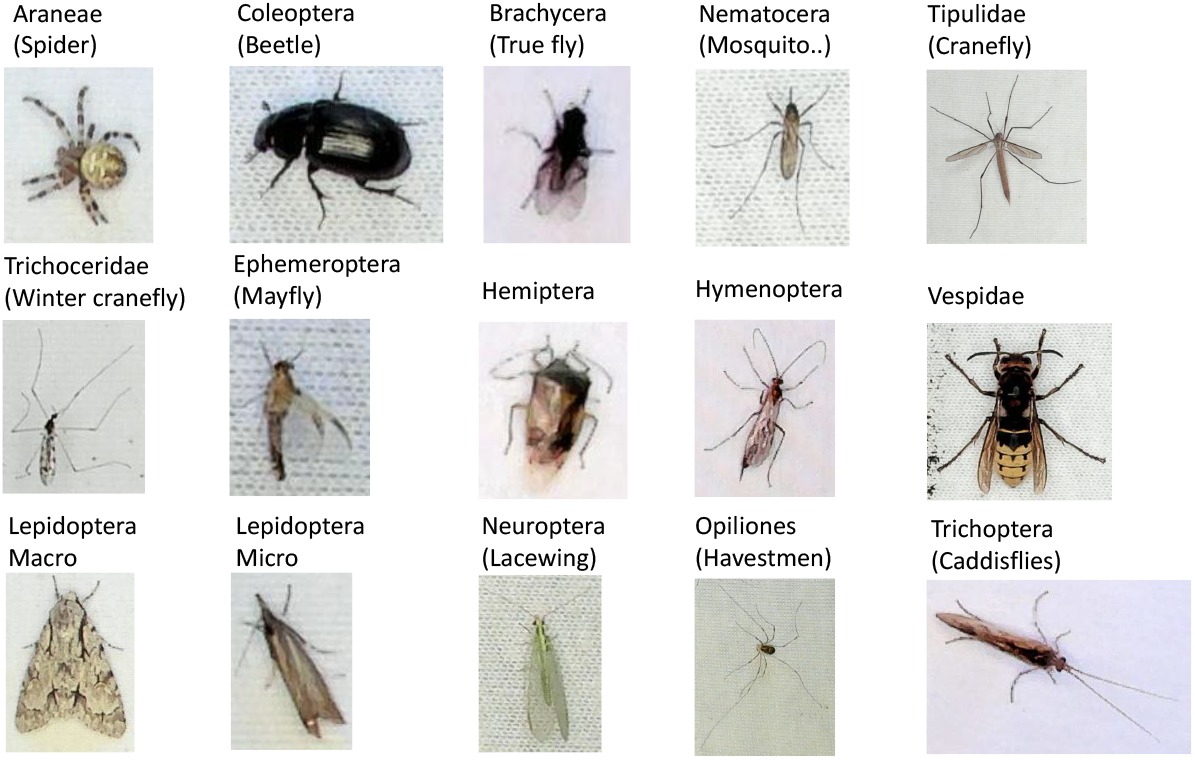
Example of samples from the 15 taxa of insect used to classify nocturnal insects in broad taxonomy ranks.

## 3 RESULTS

### 3.1 Insect detection and localization

The precision, recall and F1 score of the first stage of the pipeline is listed in Table 3. It is observed that the large YOLOv5m6 model has the highest F1 score; however, the number of parameters for YOLOv5m6 is 35.7M compared to 12.6M for YOLOv5s6. Experiments with newer models, such as YOLOv8m, did not improve the F1 score, indicating that the annotated dataset needs to be refined and expanded. The discrepancy between model predictions and the annotated insects are particularly influenced by the exclusion of small insects in the annotated images.

**TABLE 3.** Validation results for insect detection and localization on dataset with 482 labels.

| Metric | YOLOv5m6 | YOLOv5s6 |
| --- | --- | --- |
| Precision | 0.923 | 0.938 |
| Recall | 0.919 | 0.886 |
| F1-score | 0.921 | 0.911 |

### 3.2 Broad taxon classifier with anomaly detection

The precision, recall and F1 score of the second stage of the pipeline are summarized in Table 4. The table shows the results for the ResNet50v2 model evaluated without the anomaly threshold detector and where the uncertain samples are removed. There is a small increase in all metrics, which indicates that removing samples predicted as “uncertain” improves the classifier by accepting that 1.0% of the true positive samples are ignored.

**TABLE 4.** Validation metrics for the broad taxon classifier. ResNet50v2 without uncertain are the metrics where 235 samples are removed by the anomaly threshold detector.

|  | ResNet50v2 | ResNet50v2<br>without uncertain samples |
| --- | --- | --- |
| Samples | 25,034 | 24,799 |
| Precision | 0.974 | 0.977 |
| Recall | 0.965 | 0.971 |
| F1-score | 0.970 | 0.974 |

The confusion matrix for the broad taxon classifier is shown in Figure 5. Here, we have included the uncertain class for predictions below the anomaly threshold. High values are observed in the diagonal of the matrix, indicating an accurate classification. However, difficulties are observed in classifying winter craneflies (Diptera Trichoceridae) from mosquitoes (Diptera Nematocera) and craneflies (Diptera, Tipulidae); this is due to a visually similar appearance and possible errors in the training dataset for these families of Diptera.

**FIGURE 5.**
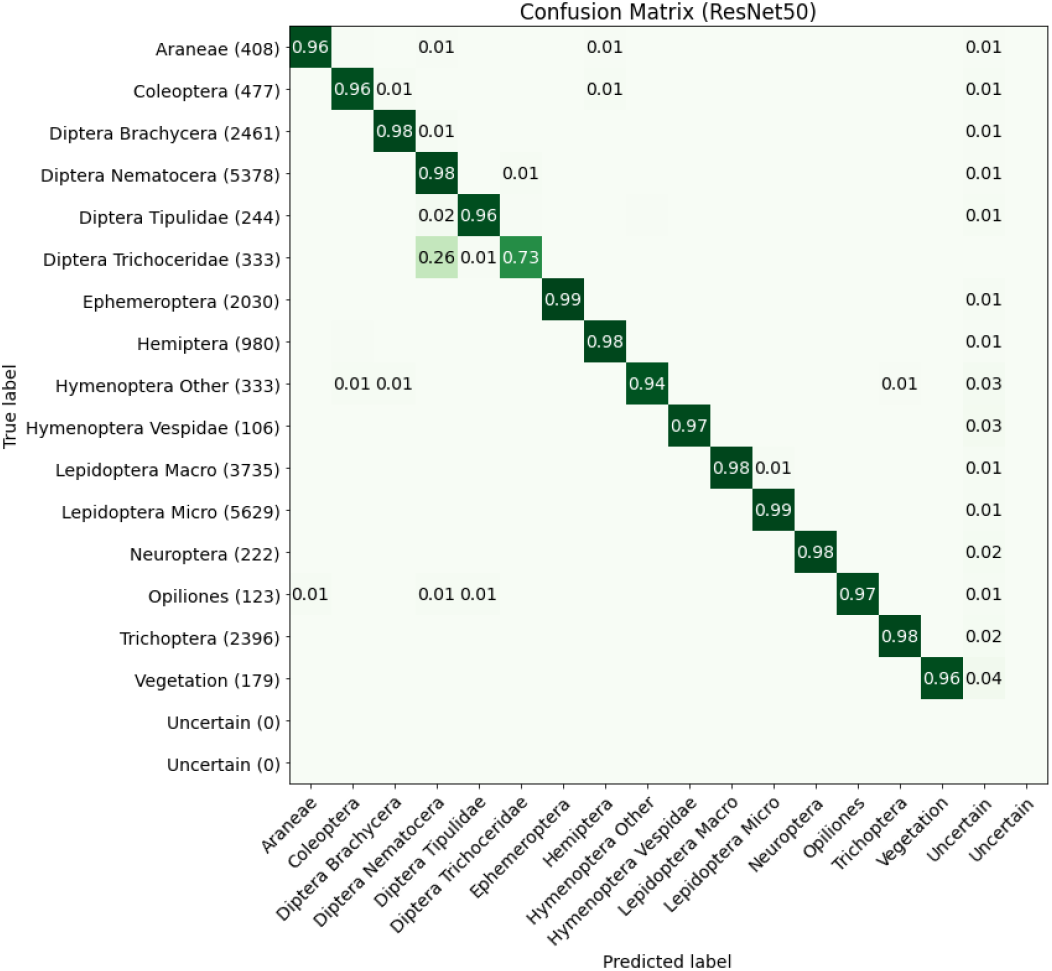
Confusion matrix for the broad taxon classifier with out-of-distribution detection of samples marked as the uncertain class.

We evaluated the broad taxon classifier with anomaly detection in the 10-minute time-lapse recordings from 2022 and 2023. This was done by selecting up to 200 randomly classified insects of the 16 taxa above and below the threshold learned from the output distribution of the dataset. We manually verified by visual inspection of the classified insects above and below the anomaly threshold. The results listed in Table 5 show that in total, 92. 3% of the insects are classified above the threshold with a precision of 96.8%. The remaining 7.7% insect detections classified as uncertain below the anomaly threshold have a similarly high precision of 95.9%. Spiders (Araneae) are the group of animals with the lowest precision of 83% above the threshold. This is because many of the false positive detections are spider webs or dirt since the training data do contain spiders with prey and more blurry and unclear objects.

**TABLE 5.** Number of classified insects taxa above and below anomaly threshold (TH) with precision based on maximum 200 random selected predictions. Above precision is the precision for classified insect taxa. Below precision is the precision for uncertain classified insects.

| Taxon category | Above TH | Below TH | Above (%) | Above precision (%) | Below precision (%) |
| --- | --- | --- | --- | --- | --- |
| Araneae | 4,957 | 864 | 85 | 83 | 100 |
| Coleoptera | 2,896 | 351 | 89 | 96 | 98 |
| Diptera Brachycera | 23,941 | 1,850 | 93 | 97 | 98 |
| Diptera Nematocera | 171,855 | 8,760 | 95 | 92 | 98 |
| Diptera Tipulidae | 1,858 | 56 | 97 | 98 | 100 |
| Diptera Trichoceridae | 7,070 | 26 | 100 | 100 | 96 |
| Ephemeroptera | 25,423 | 7,374 | 78 | 98 | 100 |
| Hemiptera | 8,008 | 345 | 96 | 92 | 100 |
| Hymenoptera Other | 6,906 | 182 | 97 | 100 | 92 |
| Hymenoptera Vespidae | 542 | 91 | 86 | 100 | 91 |
| Lepidoptera Macro | 46,425 | 2,871 | 94 | 100 | 89 |
| Lepidoptera Micro | 93,870 | 6,006 | 93 | 97 | 98 |
| Neuroptera | 1,094 | 60 | 95 | 100 | 95 |
| Opiliones | 673 | 50 | 93 | 97 | 98 |
| Trichoptera | 64,715 | 8,935 | 88 | 98 | 100 |
| Vegetation | 2,050 | 951 | 68 | 100 | 96 |
| Total/average | 462,283 | 38,772 | 92.3 | 96.8 | 95.9 |

### 3.3 Computational speed and power usage

The speed performance of our pipeline was evaluated across various computing platforms, including edge processing devices. These included a standard computer equipped with an Intel(R) Xeon(R) E5-2620 v4 @ 2.10GHz and an NVIDIA TITAN X Pascal GPU, as well as the NVIDIA Jetson Nano (JN) and Raspberry Pi 4 (RP4) (both with 4 GB of memory) and Raspberry Pi 5 (RP5) (with 8 GB of memory). An additional 4 GB swap file was required to execute the processing pipeline on the Jetson Nano for nights with more than 60 insects per image. This was necessary because the classification is performed in batches, processing all insects detected in one image simultaneously to enhance performance. The three edge computing devices were selected because they are roughly at the same price, with the JN being the most expensive (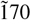 USD) at about twice the price of RP5 (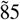 USD) since JN also has a NVIDIA Tegra X1 GPU computer. We have tested the pipeline by processing 6,271 images from one night with high insect activity. Detailed time performance and power metrics are provided in Table 6. During the test, an external SSD drive with images was connected to the Raspberry Pi and Jetson Nano computers that continuously consumed 0.9W of power.

**TABLE 6.** Processing performance on standard computer, Jetson Nano (JN) and Raspberry Pi’s (RP) of four hours recording with 6,271 images with an average of 35-37 insect detections per image.

| Platform | YOLOv5 model | Detected insects | Detector sec/img | Classifiers ms/det. | Tracker ms/img | Average sec/img | Total hours | Power (W) idle/active |
| --- | --- | --- | --- | --- | --- | --- | --- | --- |
| Standard | s6 | 221,732 | 0.018 | 7.5 | 15 | 0.296 | 0.52 | 40/90 |
| Standard | m6 | 231,835 | 0.032 | 6.7 | 15 | 0.295 | 0.51 | 40/90 |
| JN | s6 | 221,732 | 0.351 | 57.6 | 197 | 2.58 | 4.50 | 3.7/10.2 |
| JN | m6 | 231,835 | 0.848 | 58.8 | 197 | 3.22 | 5.61 | 3.7/10.2 |
| RP5 | s6 | 221,732 | 1.97 | 193 | 88 | 8.87 | 15.45 | 3.5/10.4 |
| RP5 | m6 | 231,835 | 4.30 | 189 | 88 | 11.4 | 19.82 | 3.5/10.4 |
| RP4 | s6 | 221,732 | 5.30 | 464 | 169 | 21.9 | 38.14 | 3.0/6.8 |
| RP4 | m6 | 231,835 | 9.99 | 484 | 169 | 28.1 | 48.87 | 3.0/6.8 |

### 3.4 Recording statistics

12 traps were in operation for a total of 1,399 nights during the first year of 2022, where 3,428,430 images were captured with 45,285,526 insect detections and 94% were above the anomaly threshold and used for time-lapse analysis. 97% of the detections contribute to 1,177,728 valid insect tracks with a duration of more than 4 seconds and a classification certainty of >50% for the classified insect. A summary of detailed statistics for all traps in 2022 is found in the Appendix Table A1 and for statistics in 2023 see Appendix Table A2.

Additionally, three edge-processing camera systems (RP4 and JN) have demonstrated promising reliability during monitoring with upload of abundance statistics for three months in 2024.

### 3.5 Tracking and time-lapse sampling

We hypothesize that tracking with high time-lapse sampling (0.5 fps) is the most accurate method to correctly count and classify each individual insect observed in front of the camera. This method is compared to lower time-lapse sampling rates, where tracking becomes impractical for fast-moving insects. However, lower time-lapse sampling requires fewer resources for both storage and image processing.

The tracking was evaluated by creating videos with insect classification and colored tracks, as illustrated in the appendix figure A1. Video sequences with many fast-moving insects of similar classes did have some difficulties, see example video^*§*^ with moderate insect activity. Here, it was observed that an insect track will occasionally be associated with the wrong insect. This problem could be minimized by lowering the cost threshold, but it requires a higher sampling rate to ensure that the distance an insect has moved between two frames is short.

An evaluation was made by comparing insect abundance using insect tracking in images recorded with 0.5 fps compared to selecting time-lapse images recorded by different time intervals. For each trap and taxonomic group, the abundance was calculated using tracking and the result was compared with a time-lapse approach. The time-lapse approach did not include tracking since large sample intervals were used. Here, we used localization and classification with anomaly detection by removing detections with a classification score below the learned threshold described in Section 2.4.1. We have simulated time-lapse sampling of 10s, 30s, 1m, 2m, 5m, 10m, 15m, 20m and 30m by only including image detections with these intervals.

The mean absolute difference between the time series of tracks and the time-lapse detections is shown in Figure 6. The red lines show the average difference for all traps for each insect category. We assume that the minimum difference would be the optimal time-lapse sampling interval approximating the numbers obtained by tracking. For short TL intervals (below minimum), the absolute difference is high since there are more detections than tracks. For longer TL intervals (above minimum), the absolute difference increases again, which indicates that more tracks than detections are encountered.

**FIGURE 6.**
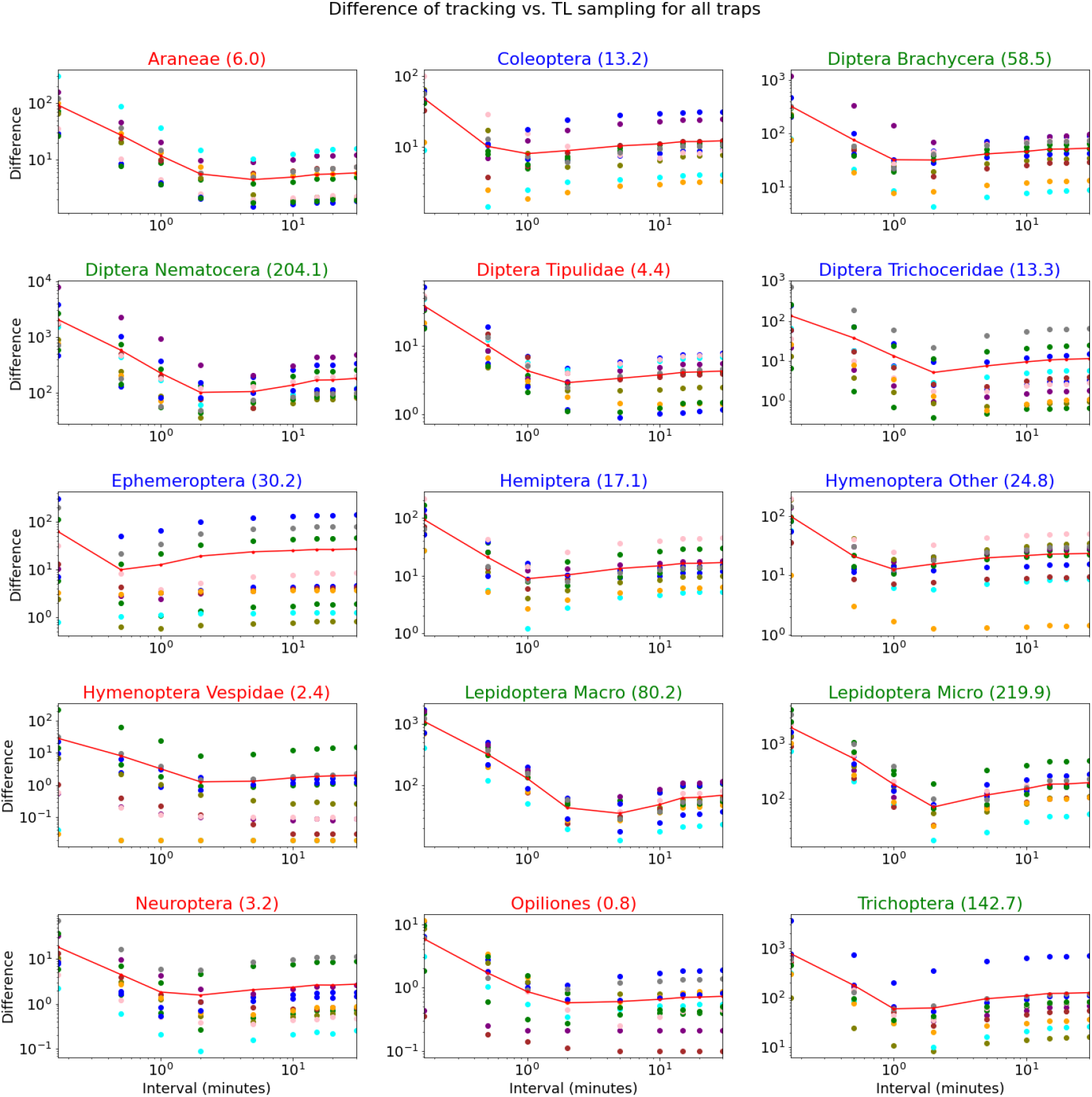
Shows the mean absolute difference between tracks and time-lapse (TL) intervals (10s-30m) for all nocturnal insects and traps (Difference marked with “•” of different colors for each trap). The average difference for all traps is shown with a red line. The values in brackets are the average number of tracks per night. Green indicates taxa with >50 tracks/night. Blue indicates taxa with 10-50 tracks/night and red indicates taxa with <10 tracks/night.

Pearson’s correlation was also applied to time series of insect tracks and time-lapse insect detections to compare the temporal dynamics of the two metrics of activity. An example of the correlation among four different traps and insect groups is shown in Appendix Figures A2 and A3. The highest correlation is achieved by 5-minute sampling intervals except for Vespidae, where the best correlation is achieved with 10-second intervals. The same tendency is observed for all time-lapse sampling intervals compared to tracking, although the number of observed detections and tracks differs.

In Appendix Figure A4 we have summarized the Pearson correlation of time-lapse sampling intervals and tracks for traps and the 15 broad arthropod taxa. The correlation is generally high (0.9) for abundant taxa with >50 tracks/night. For taxa with abundance <10 tracks/night, the correlation drops for sampling intervals greater than 10 minutes.

The duration of each track should also be considered when deciding on a time-lapse interval. Appendix Figure A5 shows the distribution of track duration for each of the 15 taxonomic groups from the camera trap with high activity of insects. The average duration ranges from 29 to 115 seconds and varies among the different taxa. In particular, lepidoptera have a longer track duration than most other groups.

Figure 7 summarizes the frequency of minimum difference and gives the best correlation of sample intervals with tracking. It indicates that a time-lapse interval of 2 minutes achieves in more than 60 situations (combinations of traps and arthropod taxa) the minimum difference between time-lapse detections and tracking. The best Pearson correlation is achieved for 10 minutes; however, 5 and 2-minute TL intervals are also a good choice as an alternative to tracking with 0.5fps. This approach would reduce the number of recorded images and allow one to perform edge processing of images in real-time on the camera trap with Raspberry Pi.

**FIGURE 7.**
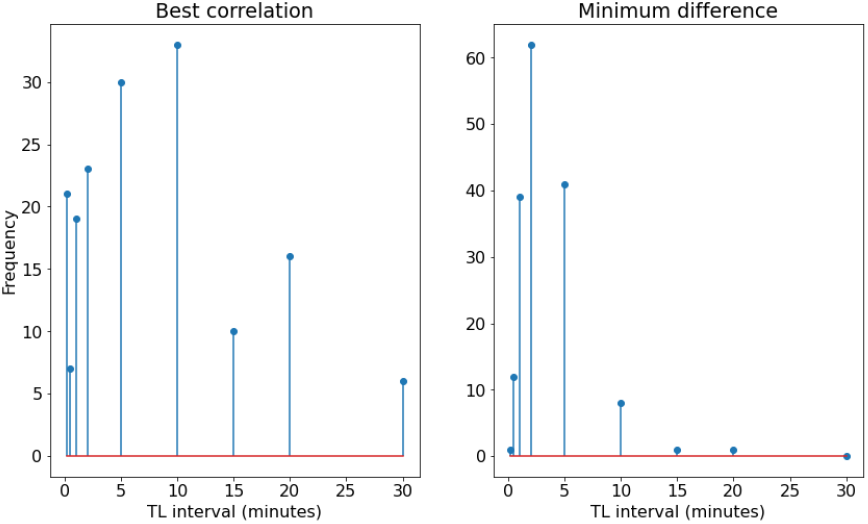
Show the frequency of best correlation and minimum difference between tracking (0.5fps) and different time-lapse intervals. Each observation concerns the best correlation or minimum difference for one trap and one of the 15 broad arthropod taxa.

### 3.6 Monitoring insect abundance

The image processing pipeline allows for the extraction of rich ecological information such as indicators of the phenology, relative abundance, and richness of insect and arachnids from insect camera trap images. This section presents an example of ecologically relevant data from one of the insect camera traps with high insect activity (LV2) deployed in Lille Vildmose during the 2023 growing season. Figure 8 shows the abundance of arthropods tracked and categorized by the broad taxon classifier. The seasonal dynamics of each taxa is characterized by activity during a large part of the season with strong day-to-day variation in number of tracks. The seasonal dynamics of the moth species with the largest number of tracks is shown in 9. Compared to the broad taxonomic groups, detections of individual species are confined to a smaller and species-specific part of the season. Similar to the dynamics of the broad taxonomic groups, the number of tracks of individual moth species vary substantially from day to day. Figure 10 shows that some species of moths dominate the Lepidoptera community. A total of 635 distinct moth species were classified, with 360 species having more than five recorded observations across 12 traps and two seasons.

**FIGURE 8.**
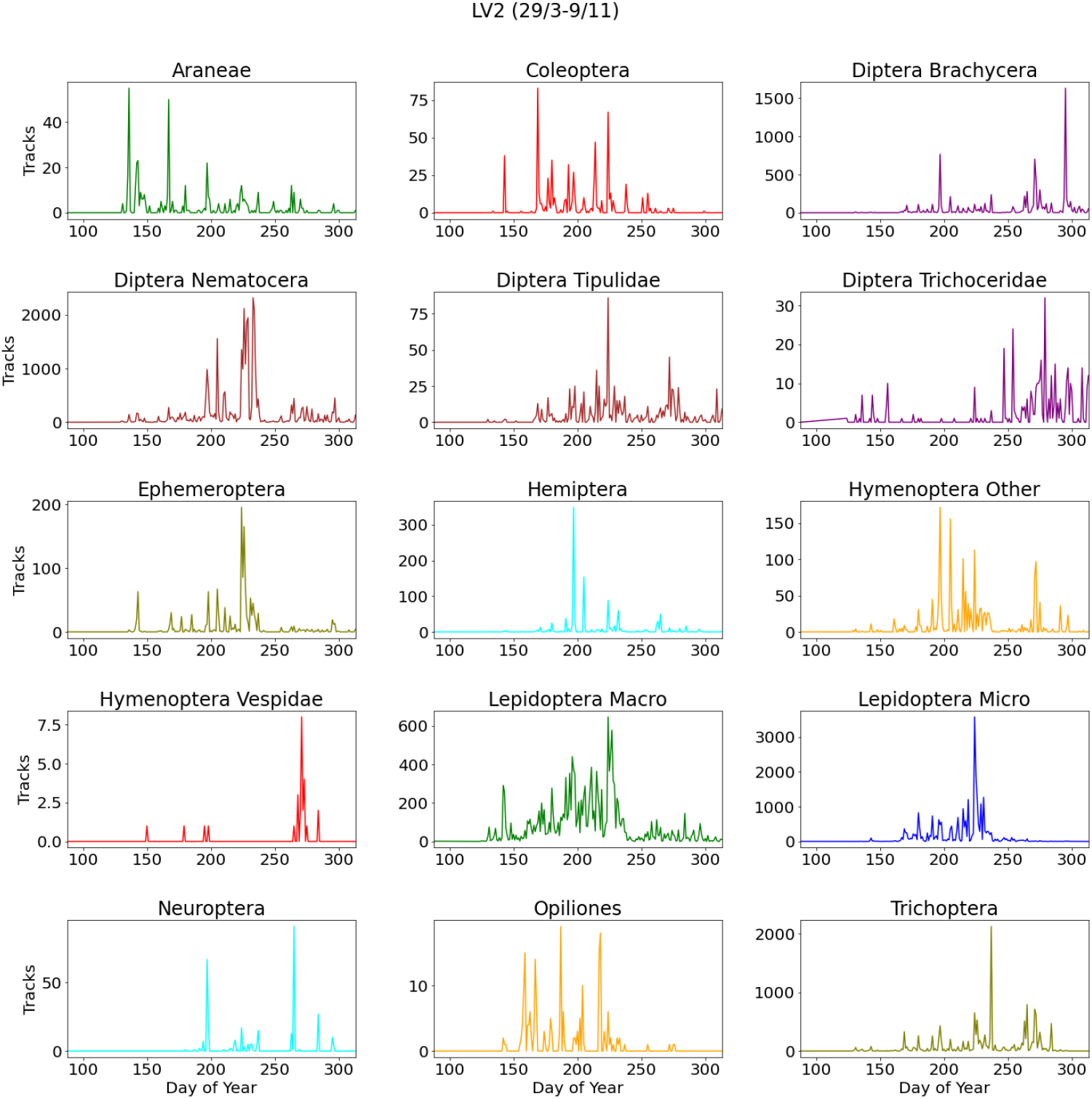
Show the abundance of the 15 insect groups of broad taxonomy observed by trap LV2 in 2023.

**FIGURE 9.**
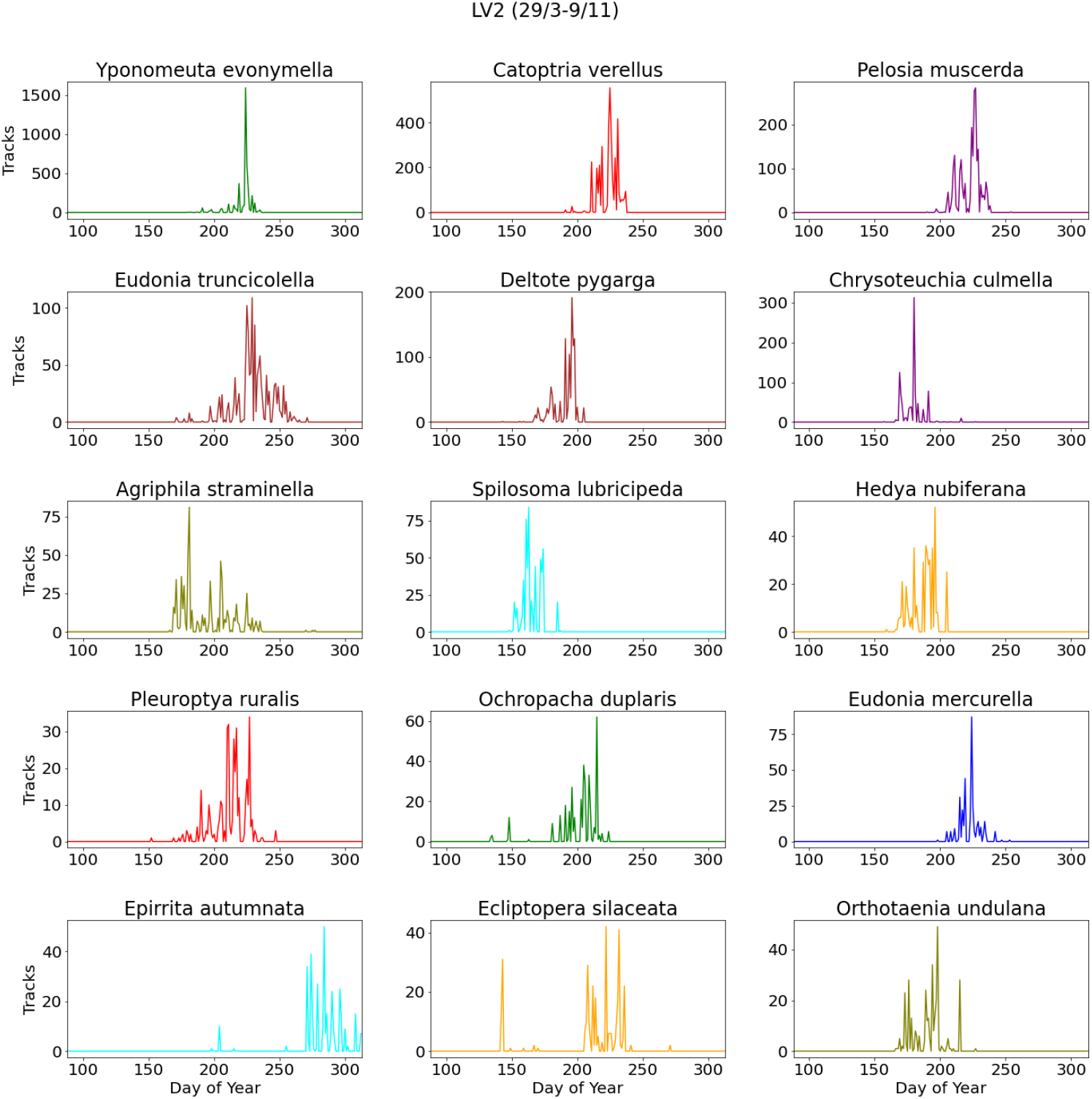
Show the abundance of 15 moths species with the highest abundance observed by trap LV2 in 2023.

**FIGURE 10.**
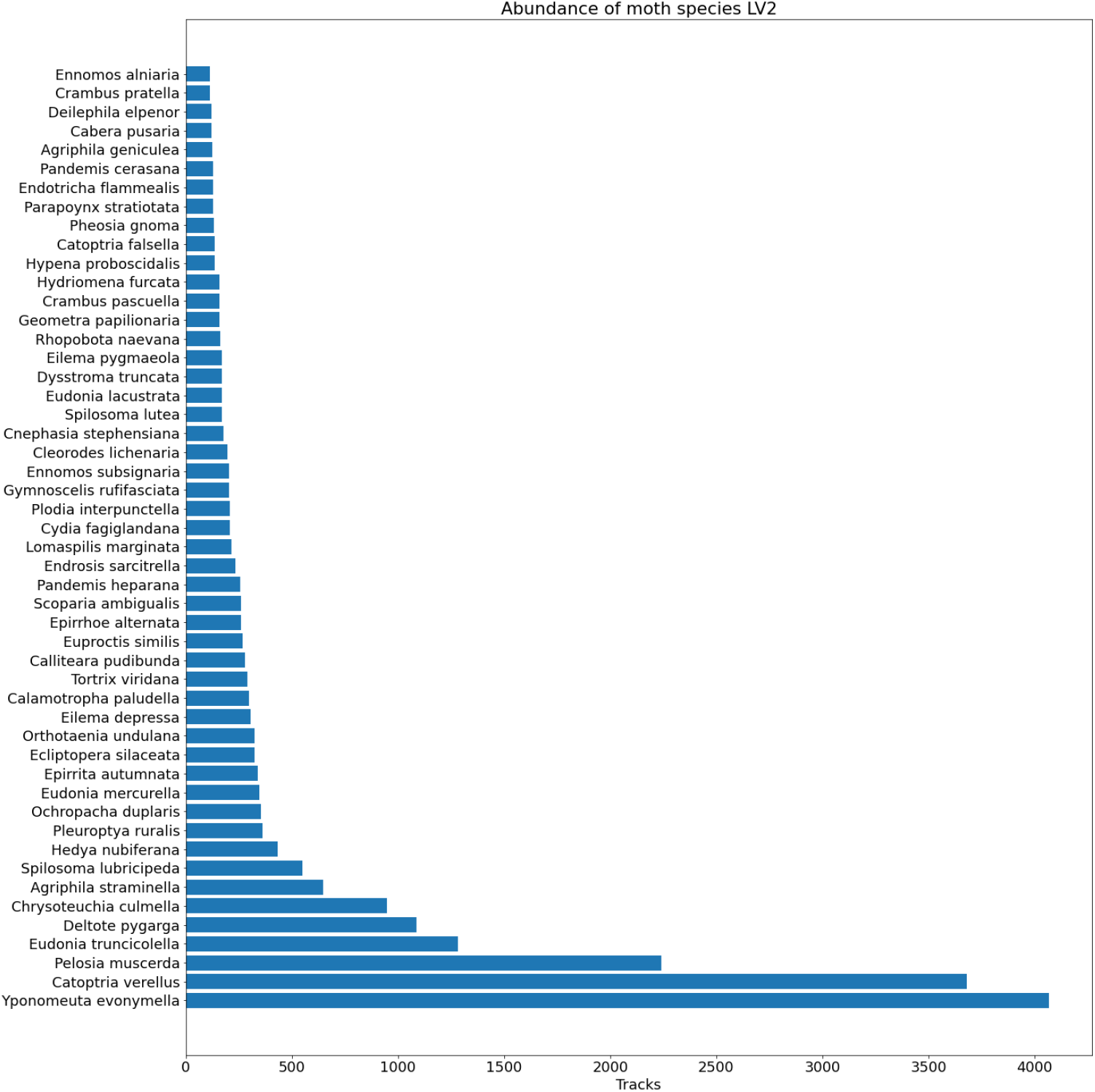
Show the most dominant moth species observed by trap LV2 in 2023.

## 4 DISCUSSION

This work proposes a novel deep learning pipeline for monitoring nocturnal insects and arachnids. The pipeline performs detection, classification and tracking within selected taxonomic groups, as well as filtering anomalies such as blurry, dark, partly visible, or uncertain insect images. In particular, it features a new image classifier to separate insects from insect camera traps into broad taxonomic groups. We have shown that the pipeline can run on edge platforms and have demonstrated its application on images recorded with 12 insect camera traps installed in bogs, heaths and forests across two full seasons.

Our pipeline features several improvements compared to previous studies. First, our YOLOv5m6 object detector model exhibits a 4.4% higher detection rate than YOLOv5s6 similar to what was found in (Bjerge et al. 2023a). However, while YOLOv5s6 shows a slightly lower recall during evaluation, it boasts faster processing speeds, particularly evident on Raspberry Pi devices. Surprisingly, the performance advantage observed with YOLOv5s6 on Raspberry Pi platforms contrasts with the less significant improvement noted when utilizing GPU acceleration.

Second, the broad taxon classifier with anomaly detection achieved a high precision of 95.9% when evaluated on time-lapse recordings taken at 10-minute intervals. We identified 7.7% anomalies from 462,283 insect detections, indicating that our anomaly filter effectively removes uncertain classifications. The classification of insects and arachnids into 16 broad taxonomic groups achieved a precision of 96.8%, based on an evaluation of 200 random trap images. These results indicate that the anomaly detector finds and removes uncertain anomalies such as debris, partially visible or blurry insects, or insects belonging to groups not represented in the training data.

Third, we developed a simple track-by-detection algorithm that matches feature embeddings, distance and area for insect tracking in time lapse images at a frame rate of 0.5 fps. The tracking algorithm has been preliminarily evaluated by visually inspecting the videos created by the proposed pipeline. However, the same algorithm, but without the cost of embedding feature, was already evaluated by Bjerge et al. (2021b). Even if the amount of data collected is substantial, the frame rate is rather low for tracking and can sometimes be inaccurate for moving insects of the same species due to the long distances they can travel between frames. More work is needed to improve the tracking algorithm before conducting a thorough evaluation.

In addition to the AMI moth species classifier (Jain et al. 2025), a growing number of classification models are available. For example, the European moth species classifier has high accuracy in high-quality images and is trained on an extensive dataset in terms of species and images (Korsch, D., Bodesheim, P., Denzler, J. 2021). Other models capable of classifying moth species from specific regions such as North-Eastern North America (Quebec/Vermont) and Panama are also available (Rolnick, David; Bundsen, Michael; Jain, Aditya; Cunha 2023). In other studies a single model for classification with hierarchical taxonomic ranks has been proposed (Bjerge et al. 2023b). However, creating a dataset and code that enable the training and evaluation of such a model requires further work.

We compared tracking with a standard time-lapse sampling approach without tracking. Lower time-lapse sampling intervals (10 seconds to 30 minutes) significantly reduce the amount of data collected. We found a high correlation between tracking and time-lapse sampling, particularly for taxa with more than 50 tracks per night. For taxa with fewer than 10 tracks per night, the correlation decreases for time-lapse intervals longer than 10 minutes. The best mean absolute difference was found for 2-minute intervals, but for taxa with fewer than one detection per night, the correlation decreases.

Edge processing on the JN platform is by far the fastest and most power-efficient since RP5 and JN nearly consume the same amount of power, but JN is more than four times faster processing image with an average of 2.6 seconds. RP4 is the slowest platform. However, the average processing (28.1s) time is still less than 30 seconds and, therefore, still suitable for time-lapse sampling down to intervals of 30 seconds. when processing in real-time is performed. The most power-efficient processing platform is the JN, which on average consumes 26.3 Joule per image (YOLOv5s6) where RP4 consumes 148.9 Joule and RP5 92.3 Joule.

All edge processing platforms are suitable for a time-lapse sampling approach with intervals as short as 10 to 30 seconds. Among these, the JN stands out as the most energy-efficient solution, capable of real-time tracking at nearly 0.5 fps. The JN platform has also been used for the real-time monitoring of diurnal insects, as demonstrated by Bjerge et al. (2021a). Alternatively, the video monitoring platform suggested by Sittinger et al. (2024) could be considered for deploying our proposed pipeline directly on the Luxonis OAK-1 camera with processing power, although some modifications may be necessary.

Insect camera traps can generate detailed ecological indicators at the community and species level. For instance, the relative abundance and seasonal dynamics of different broad taxonomic groups of terrestrial arthropods highlight the diversity of nocturnal insects that can be monitored with a UV enabled insect camera trap. Many insect taxa, for which we have very little data, are quite abundant in the trap data, including adult stages of insects normally associated with aquatic environments. The strong day-to-day variation in activity for both broad arthropod taxa and individual moth species indicates that weather patterns play a key role in determining their occurrence in the trap (Bjerge et al. 2021b).

Since a large proportion of moth species can be identified directly from images, data from insect camera traps can describe the seasonal dynamics of an important part of the insect community. Such information could allow conservation actions to be timed according to the activity of individual species if, for instance, particular management should happen after the flight season of a particular species. Another strength of the detailed phenology information provided by the traps is the possibility of timing the search for rare species based on the phenology of related and more common species active at the same time.

The high number of species that can be detected with traps suggests that this sensor is capable of detecting even small changes in the composition of the community in response to changes in the local environment. Such changes could be the result of habitat deterioration or restoration or the result of global change drivers, including climate change.

The long-tailed distribution common to most biological communities, where few species are common and many species are rare, is also visible in insect camera trap data. This poses a challenge for the training of reliable classification models. However, given the rate at which insect camera traps collect data and how their use is increasing, we predict that even this challenge will become smaller and that there is a promising future for insect camera traps as a standardized and widespread monitoring approach.

## AUTHOR CONTRIBUTIONS

All authors have seen and approved the submitted version of the manuscript. All authors have contributed to the work and all persons entitled to coauthorship have been included. Kim Bjerge: Conceptualization, Investigation, Methodology, Coding, Formal Analysis, Writing - Original Draft, Writing — Review & Editing, Visualization. Henrik Karstoft: Supervision, Writing — Review & Editing, Supervision. Toke T. Høye: Conceptualization, Data curation, Resources, Writing — Review & Editing, Supervision, Funding.

## ACKNOWLEDGMENTS

This research was funded by Aage V. Jensen Naturfond grant number N2024-0013 and the European Union’s Horizon Europe Research and Innovation programme, under Grant Agreement No. 101060639 (MAMBO). We also acknowledge Nathan Pinoy, who has been part of collecting images and contributed to the creation of the broad taxon dataset.

## APPENDIX

### A SUPPLEMENTARY MATERIAL

**FIGURE A1.**
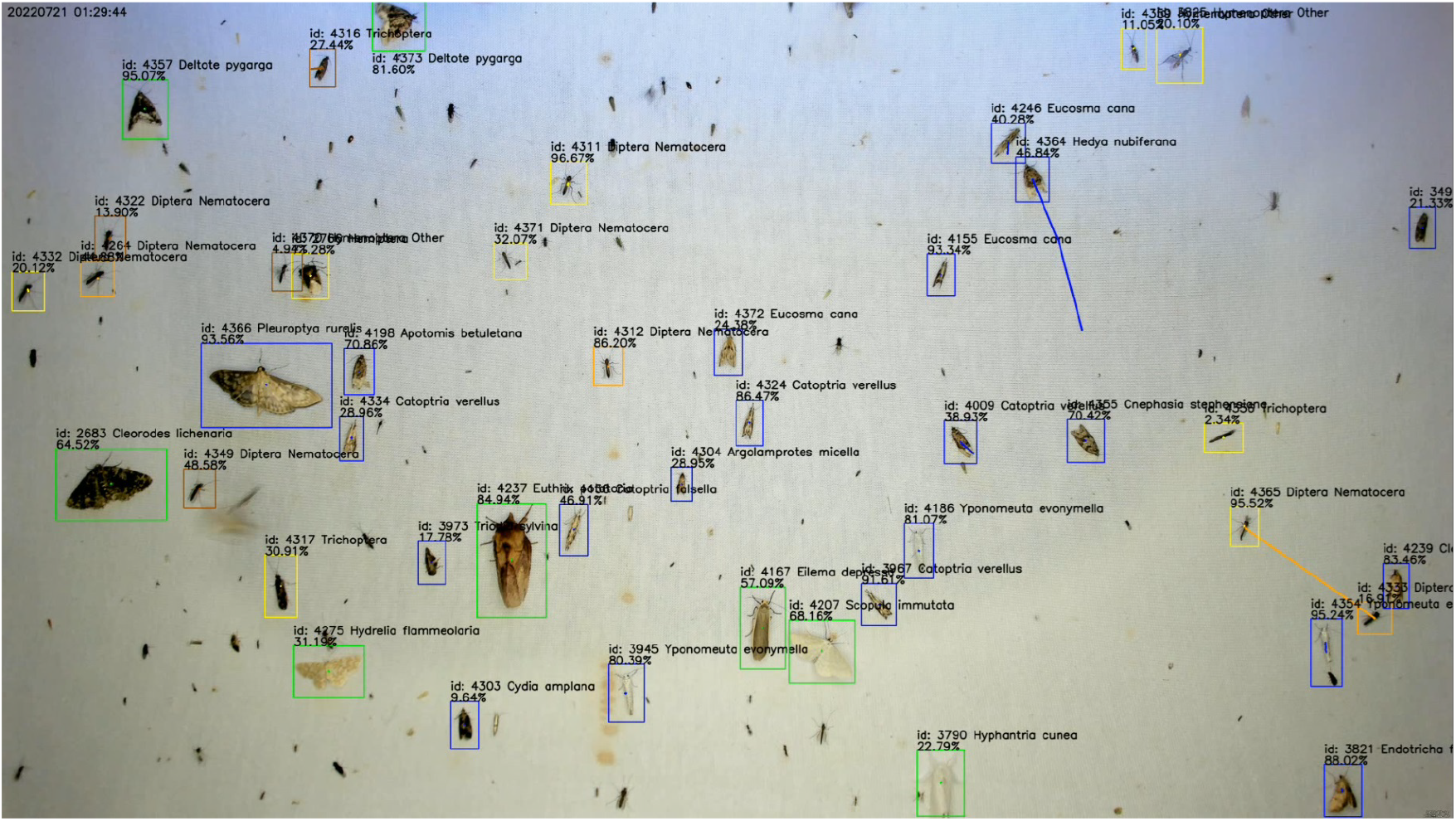
Example of image from camera trap processed with the pipeline to track and classify arthropods in 15 taxonomic groups and moth species.

Figure A1 shows an image recorded from a trap processed by the processing pipeline to detect, track and classify insects in broad taxonomic groups and moth species. Figures A2 and A3 show the abundance measured using the tracking of images recorded with 0.5 fps or filtered detections and classifications with time-lapse intervals ranging from 10 seconds to 30 minutes. The Pearson correlation between tracking and filtered classifications with varying TL intervals is shown in the title of each plot. Figure A4 shows the Pearson correlations between tracks and time-lapse intervals for all nocturnal insects and traps.

Table A1 contains list of the statistics of operational nights in 2022, number of recorded images, number of insect detections and insect tracks. Trap SS3 only collected images in the first 11 nights before tilted over by a wild boar. Trap OH1 was not operational at the start of the season due to technical issues. The power cable to trap LV1 was broken by humans driving a forest machine at the end of the monitoring season. Traps OH1-OH4 where powered by a 175W solar panel and 12V/100Ah battery. Table A2 contains list of the statistics of operational nights in 2023. Trap OH3 was not operational at the start of the season due to power supply issues. Trap SS4 missed several days of recording because animals damaged the power cable.

Figure A5 shows the histograms for the distribution of track duration for the 15 different groups of taxa from one of the traps with the highest and most diverse insect activity.

**FIGURE A2.**
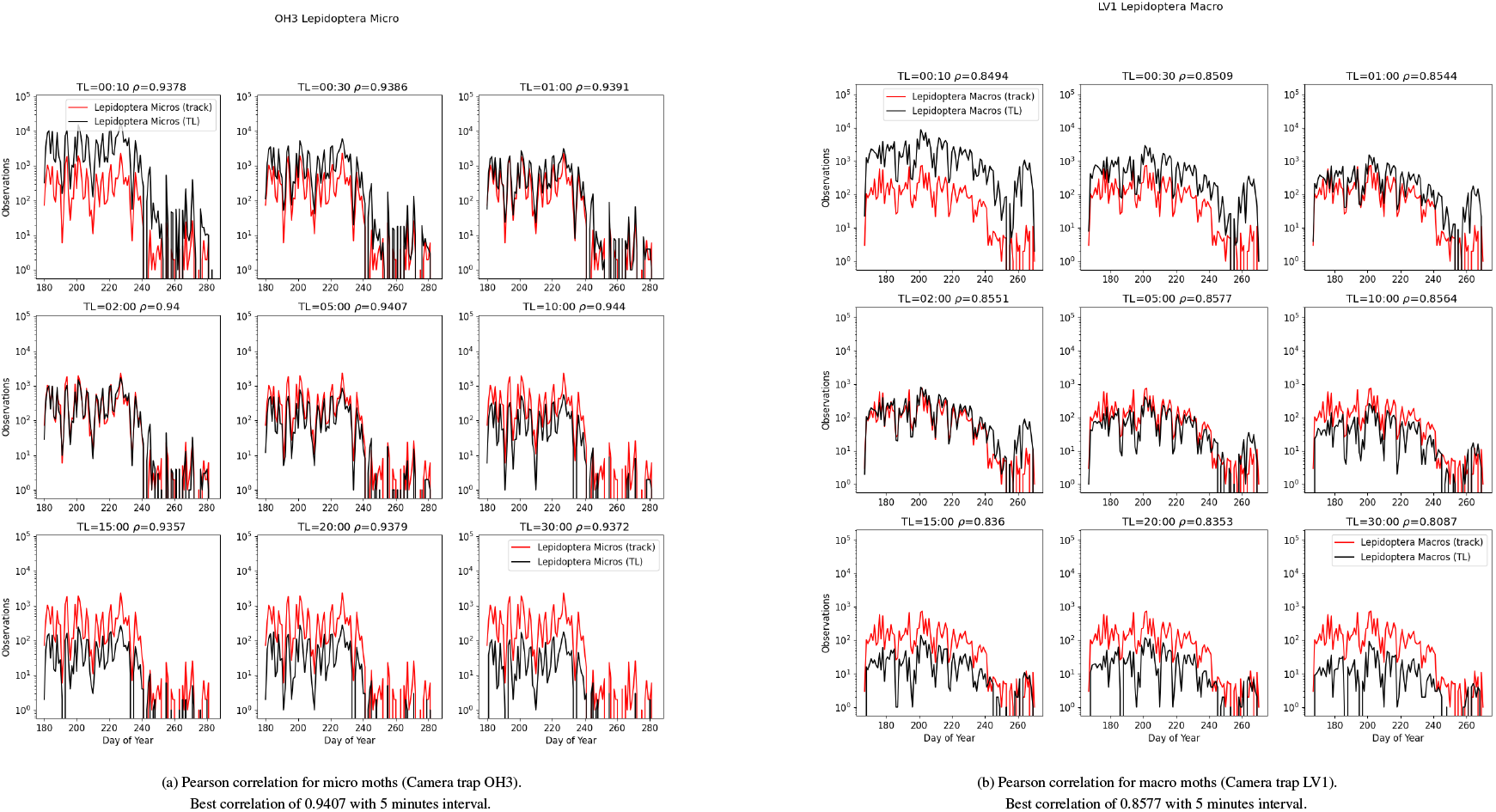
Shows the nuber of observations at several different time-lapse (TL) intervals (10s-30m) and tracking (track) for Lepidoptera micro and macro at example traps.

**FIGURE A3.**
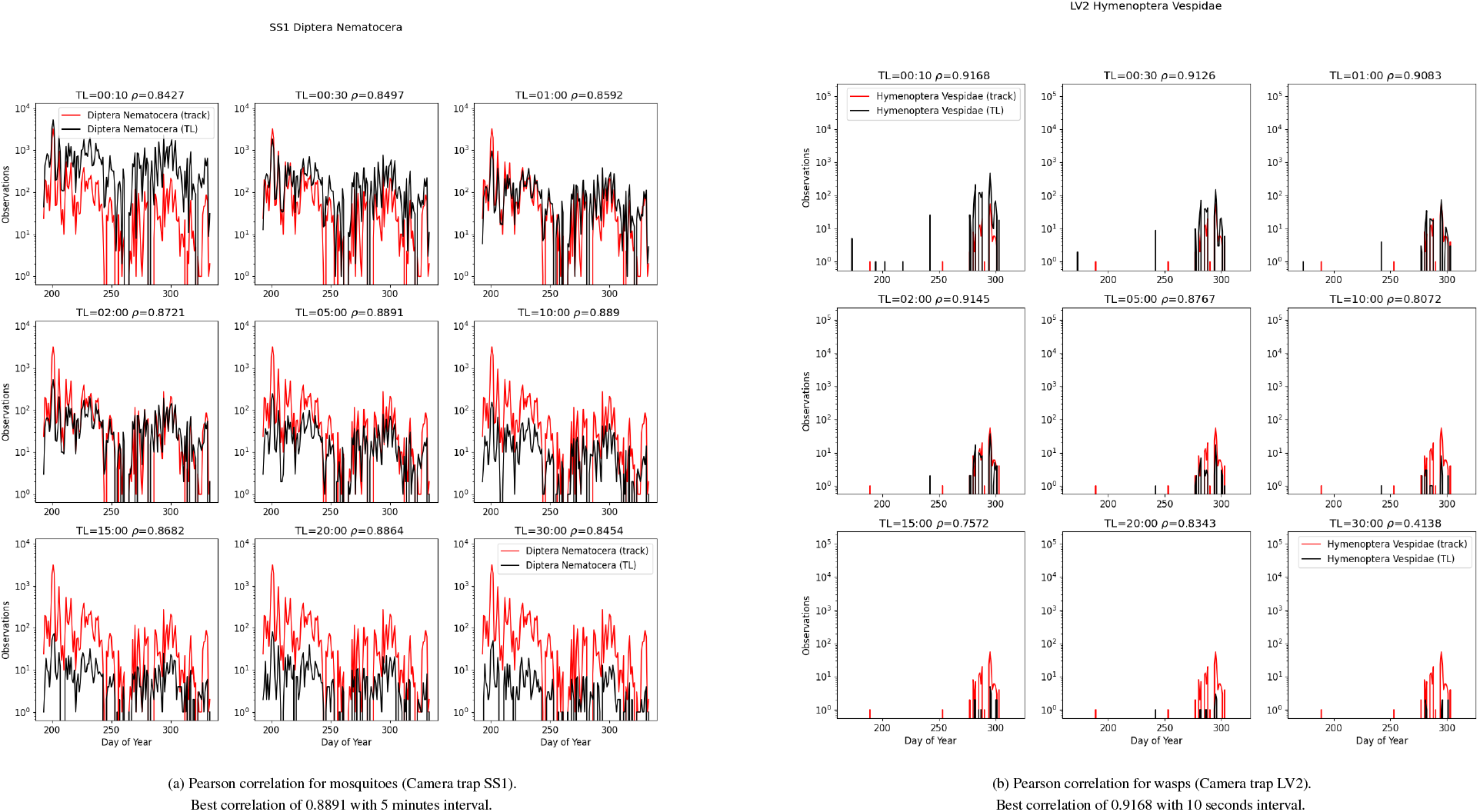
Shows the number of observations at several different time-lapse (TL) intervals (10s-30m) and tracking (track) for Diptera Nematocera and Hymenoptera Vespidae at example traps.

**FIGURE A4.**
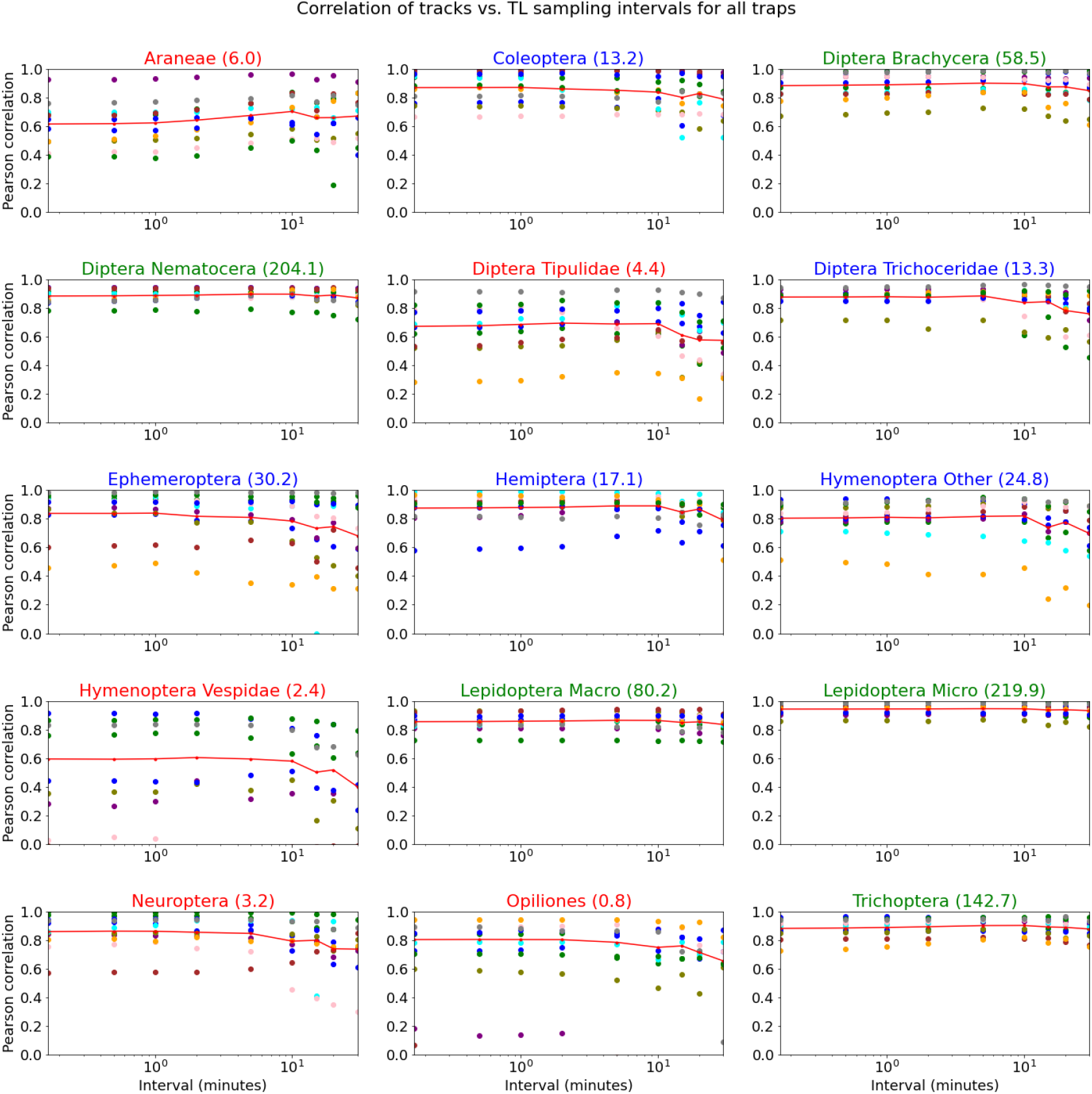
Pearson correlation between tracks and time-lapse (TL) intervals (10s-30m) for all nocturnal insects and traps (Correlation marked with “•” of different colors for each trap). The average Pearson correlation is shown with a red line. The values in parenthesis are the average number of tracks per night. Green indicates taxa with a high abundance above 50 tracks per day. Blue indicates taxa between 10-50 tracks per night and red below 10 tracks per night.

**TABLE A1.** Statistics for processing all images recorded in 2022 from 12 camera traps. The detections above the anomaly threshold (>TH) indicate the percentage of valid detections. Valid tracks are defined for insects visible more than 4 sec. and classified for the same taxa in more than 50% of the detections in a track.

| Trap | Days | Files | Detections | Detections<br>>TH (%) | Tracks | Valid<br>tracks | Detections in<br>tracks (%) | Valid<br>tracks (%) |
| --- | --- | --- | --- | --- | --- | --- | --- | --- |
| LV1 | 103 | 361,429 | 5,046,324 | 96 | 151,174 | 125,420 | 97 | 83 |
| LV2 | 151 | 464,950 | 6,742,175 | 94 | 201,241 | 167,879 | 97 | 83 |
| LV3 | 135 | 371,851 | 8,894,099 | 94 | 196,214 | 165,082 | 97 | 84 |
| LV4 | 151 | 309,317 | 2,327,043 | 94 | 76,128 | 62,495 | 97 | 82 |
| OH1 | 82 | 131,441 | 959,849 | 94 | 32,031 | 26,588 | 96 | 83 |
| OH2 | 118 | 208,048 | 2,027,460 | 94 | 66,450 | 53,052 | 96 | 80 |
| OH3 | 116 | 300,387 | 3,488,553 | 93 | 113,569 | 91,076 | 96 | 80 |
| OH4 | 111 | 246,971 | 1,387,302 | 91 | 52,089 | 41,767 | 95 | 80 |
| SS1 | 140 | 292,991 | 5,196,560 | 93 | 233,952 | 193,596 | 96 | 83 |
| SS2 | 140 | 324,401 | 33,92,246 | 95 | 115,447 | 93,216 | 96 | 81 |
| SS3 | 11 | 50,879 | 1,011,345 | 93 | 37,089 | 30,059 | 96 | 81 |
| SS4 | 141 | 365,765 | 4,809,570 | 95 | 153,823 | 127,498 | 97 | 83 |
| T/A | 1,399 | 3,428,430 | 45,282,526 | 94 | 1,429,207 | 1,177,728 | 97 | 82 |

**TABLE A2.** Statistics for processing all images recorded in 2023 from 12 camera traps. The detections above the anomaly threshold (>TH) indicate the percentage of valid detections. Valid tracks are defined for insects visible more than 4 sec. and classified for the same taxa in more than 50% of the detections in a track.

| Trap | Days | Files | Detections | Detections<br>>TH (%) | Tracks | Valid<br>tracks | Detections in<br>tracks (%) | Valid<br>tracks (%) |
| --- | --- | --- | --- | --- | --- | --- | --- | --- |
| LV1 | 169 | 226,238 | 2,912,000 | 93 | 173,203 | 141,637 | 95 | 82 |
| LV2 | 190 | 188,491 | 2,071,591 | 91 | 125,535 | 104,617 | 94 | 83 |
| LV3 | 189 | 222,269 | 3,257,249 | 93 | 170,373 | 143,401 | 95 | 84 |
| LV4 | 181 | 172,430 | 1,142,400 | 91 | 56,742 | 46,278 | 94 | 82 |
| OH1 | 200 | 177,522 | 2,961,214 | 92 | 147,120 | 121,117 | 95 | 82 |
| OH2 | 178 | 224,054 | 3,657,196 | 86 | 179,658 | 145,548 | 94 | 81 |
| OH3 | 66 | 202,204 | 1,562,553 | 89 | 74,822 | 58,072 | 94 | 78 |
| OH4 | 193 | 257,666 | 3,108,915 | 87 | 171,852 | 137,552 | 93 | 80 |
| SS1 | 125 | 165,026 | 4,869,601 | 84 | 340,321 | 278,240 | 94 | 82 |
| SS2 | 163 | 197,093 | 4,467,014 | 73 | 314,721 | 251,246 | 91 | 80 |
| SS3 | 162 | 241,957 | 2,997,824 | 71 | 204,217 | 151,454 | 89 | 74 |
| SS4 | 64 | 33,325 | 99,368 | 93 | 9,225 | 7,233 | 89 | 78 |
| T/A | 1,880 | 2,308,275 | 33,106,925 | 85 | 1,967,789 | 1,586,395 | 93 | 81 |

**FIGURE A5.**
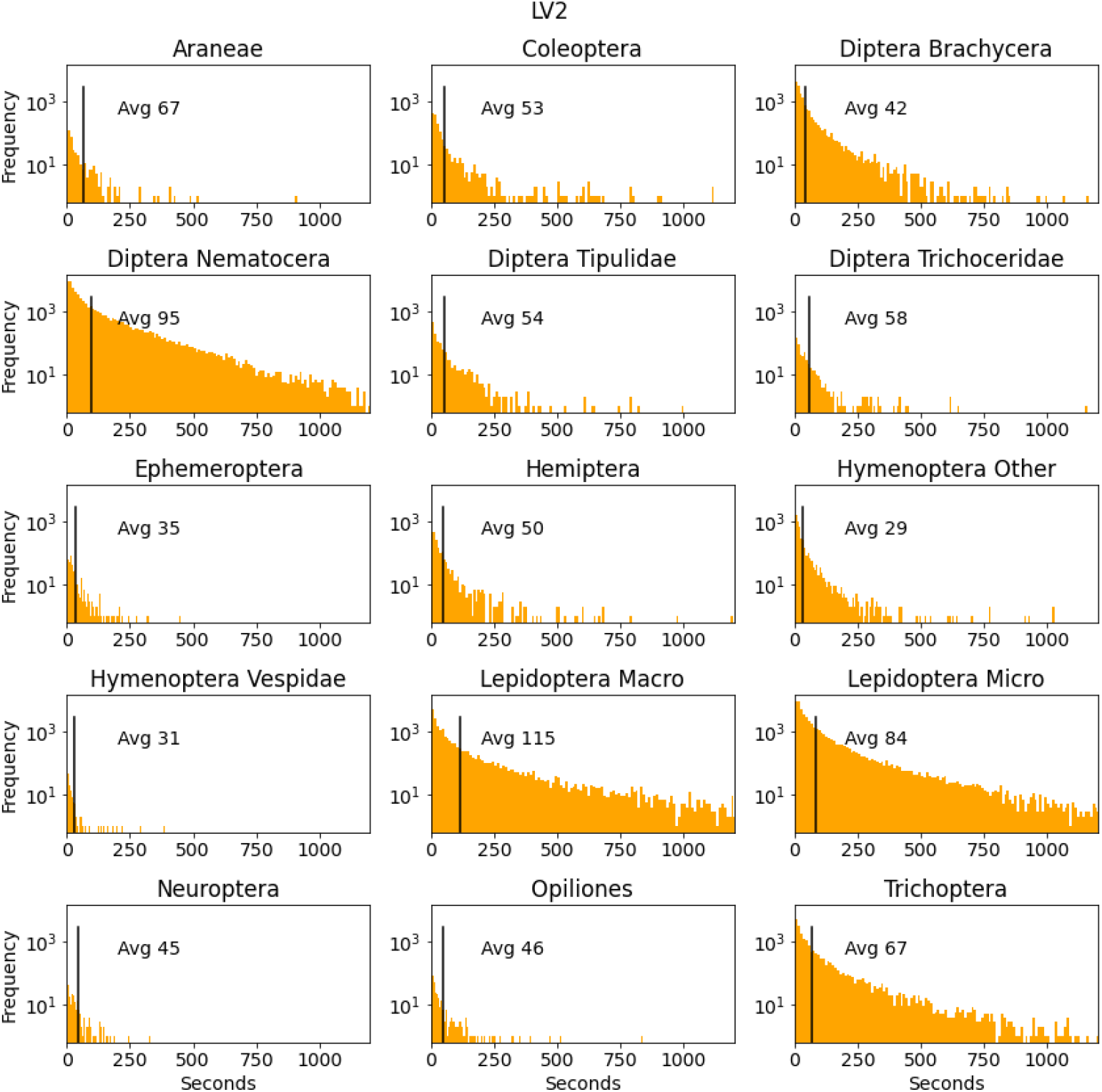
Shows the distribution of insect track durations for the 15 taxa of arthropods and the average duration of tracks in seconds from camera trap LV2. The horizontal lines show the average time of tracks in seconds. The x-axis is limited to 20 minutes, however, some insects stayed in the camera view for the whole night.

## Footnotes

† https://www.avjf.dk/avjnf/naturomraader/

‡ https://github.com/kimbjerge/MCC24-trap

§ https://www.youtube.com/watch?v=HzOCYlgnhlE

## REFERENCES

Bar-On, Y.M., Phillips, R. & Milo, R. (2018) The biomass distribution on Earth. Proceedings of the National Academy of Sciences of the United States of America, 115(25). doi:10.1073/pnas.1711842115.

Barlow, S.E. & O’Neill, M.A. (2020) Technological advances in field studies of pollinator ecology and the future of e-ecology. Current Opinion in Insect Science, 38. doi:10.1016/j.cois.2020.01.008.

Beery, S., Van Horn, G. & Perona, P. Recognition in Terra Incognita. In: Lecture Notes in Computer Science (including subseries Lecture Notes in Artificial Intelligence and Lecture Notes in Bioinformatics). Vol. 11220 LNCS, 2018.

Besson, M., Alison, J., Bjerge, K., Gorochowski, T.E., Høye, T.T., Jucker, T. et al. (2022) Towards the fully automated monitoring of ecological communities. Ecology Letters, 25(12). doi:10.1111/ele.14123.

Bjerge, K., Alison, J., Dyrmann, M., Frigaard, C.E., Mann, H.M.R. & Høye, T.T. (2023) Accurate detection and identification of insects from camera trap images with deep learning. PLOS Sustainability and Transformation, 2(3). doi:10.1371/journal.pstr.0000051. URL https://doi.org/10.1371/journal.pstr.0000051

Bjerge, K., Geissmann, Q., Alison, J., Mann, H.M.R., Høye, T.T. & Dyrmann, M. (2023) Hierarchical Classification of Insects with Multitask Learning and Anomaly Detection. Ecological Informatics, 77(102278). doi:10.1016/j.ecoinf.2023.102278.

Bjerge, K., Mann, H.M. & Høye, T.T. (2021) Real-time insect tracking and monitoring with computer vision and deep learning. Remote Sensing in Ecology and Conservation, 8(6). doi:10.1002/rse2.245.

Bjerge, K., Nielsen, J.B., Sepstrup, M.V., Helsing-Nielsen, F. & Høye, T.T. (2021) An automated light trap to monitor moths (Lepidoptera) using computer vision-based tracking and deep learning. Sensors (Switzerland), 21(2). doi:10.3390/s21020343.

Bulusu, S., Kailkhura, B., Li, B., Varshney, P.K. & Song, D. (2020) Anomalous Example Detection in Deep Learning: A Survey. IEEE Access, 8. doi:10.1109/ACCESS.2020.3010274.

Didham, R.K., Basset, Y., Collins, C.M., Leather, S.R., Littlewood, N.A., Menz, M.H. et al. (2020) Interpreting insect declines: seven challenges and a way forward. Insect Conservation and Diversity, 13(2). doi:10.1111/icad.12408.

Geissmann, Q., Abram, P.K., Wu, D., Haney, C.H. & Carrillo, J. (2022) Sticky Pi is a high-frequency smart trap that enables the study of insect circadian activity under natural conditions. PLoS Biology, 20(7), 1–26. doi:10.1371/journal.pbio.3001689.

Glenn Jocher (2020) You Only Look Once Ver. 5 (YOLOv5) on Github. https://github.com/ultralytics/yolov5 [Accessed: 21.05.2024].

He, K., Zhang, X., Ren, S. & Sun, J. Deep residual learning for image recognition. In: Proceedings of the IEEE Computer Society Conference on Computer Vision and Pattern Recognition. Vol. 2016-Decem, 2016.

Høye, T.T., Ärje, J., Bjerge, K., Hansen, O.L.P., Iosifidis, A., Leese, F. et al. (2021) Deep learning and computer vision will transform entomology. Proceedings of the National Academy of Sciences, 118(2). doi:10.1073/pnas.2002545117.

Jain, A., Cunha, F., Bunsen, M.J., Cañas, J.S., Pasi, L., Pinoy, N. et al. Insect identification in the wild: The ami dataset. In: Leonardis, A., Ricci, E., Roth, S., Russakovsky, O., Sattler, T. & Varol, G. (Eds.) Computer Vision – ECCV 2024, 2025. Cham: Springer Nature Switzerland, pp. 55–73.

Korsch, D., Bodesheim, P., Denzler, J. (2021) Deep Learning Pipeline for Automated Visual Moth Monitoring: Insect Localization and Species Classification. INFORMATIK 2021. Gesellschaft für Informatik, Bonn.,, 443–460. doi:10.18420/informatik2021-036. URL https://dl.gi.de/handle/20.500.12116/37700

Lima, M.C.F., Leandro, M.E.D.d.A., Valero, C., Coronel, L.C.P. & Bazzo, C.O.G. (2020) Automatic detection and monitoring of insect pests - A review. Agriculture (Switzerland), 10(5). doi:10.3390/agriculture10050161.

Lin, T.Y., Maire, M., Belongie, S., Bourdev, L., Girshick, R., Hays, J. et al. (2015) Microsoft COCO: Common Objects in Context. Proceedings of the IEEE Computer Society Conference on Computer Vision and Pattern Recognition,.

Logitech (2021) Brio ultra hd pro webcam. https://www.logitech.com/en-us/products/webcams/brio-4k-hdr-webcam.960-001105.html [Accessed: 21.05.2024].

Möglich, J.M., Lampe, P., Fickus, M., Younis, S., Gottwald, J., Nauss, T. et al. (2023) Towards reliable estimates of abundance trends using automated non-lethal moth traps. Insect Conservation and Diversity,(April 2022), 1–11. doi:10.1111/icad.12662.

Montgomery, G.A., Belitz, M.W., Guralnick, R.P. & Tingley, M.W. (2021) Standards and Best Practices for Monitoring and Benchmarking Insects. Frontiers in Ecology and Evolution, 8. doi:10.3389/fevo.2020.579193.

Motion (2021) Motion an open source program that monitors video from cameras. https://motion-project.github.io/ [Accessed: 21.05.2024].

Oliver, R.Y., Iannarilli, F., Ahumada, J., Fegraus, E., Flores, N., Kays, R. et al. (2023) Camera trapping expands the view into global biodiversity and its change. Philosophical Transactions of the Royal Society B: Biological Sciences, 378(1881). doi:10.1098/rstb.2022.0232.

Ratnayake, M.N., Dyer, A.G. & Dorin, A. (2021) Tracking individual honeybees among wildflower clusters with computer vision-facilitated pollinator monitoring. PLOS ONE, 16(2), 1–20. doi:10.1371/journal.pone.0239504. URL https://doi.org/10.1371/journal.pone.0239504

Redmon, J., Divvala, S., Girshick, R. & Farhadi, A. You only look once: Unified, real-time object detection. In: Proceedings of the IEEE Computer Society Conference on Computer Vision and Pattern Recognition, 2016.

Rolnick, David; Bundsen, Michael; Jain, Aditya; Cunha, F. (2023) AMI data companion. https://github.com/RolnickLab/ami-data-companion [Accessed: 21.05.2024].

Roy, D.B., Alison, J., August, T.A., Bélisle, M., Bjerge, K., Bowden, J.J. et al. (2024) Towards a standardized framework for of AI-assisted, image-based monitoring of nocturnal insects. Philosophical Transactions B, 379(20230108). doi:10.1098/rstb.2023.0108.

Russakovsky, O., Deng, J., Su, H., Krause, J., Satheesh, S., Ma, S. et al. (2015) ImageNet Large Scale Visual Recognition Challenge. International Journal of Computer Vision, 115(3). doi:10.1007/s11263-015-0816-y.

Sittinger, M., Uhler, J., Pink, M. & Herz, A. (2024) Insect detect: An open-source DIY camera trap for automated insect monitoring. PLoS ONE, 19(4 April), 1–28. doi:10.1371/journal.pone.0295474. URL http://dx.doi.org/10.1371/journal.pone.0295474

van Klink, R., August, T., Bas, Y., Bodesheim, P., Bonn, A., Fossøy, F. et al. (2022) Emerging technologies revolutionise insect ecology and monitoring. Trends in Ecology and Evolution, 37(10), 872–885. doi:10.1016/j.tree.2022.06.001.

Wagner, D.L., Grames, E.M., Forister, M.L., Berenbaum, M.R. & Stopak, D. (2021) Insect decline in the Anthropocene: Death by a thousand cuts. Proceedings of the National Academy of Sciences of the United States of America, 118(2). doi:10.1073/PNAS.2023989118.

